# Time-dependent modulation of gut microbiome in response to systemic antifungal agents

**DOI:** 10.1101/2020.10.05.315184

**Authors:** Nadeeka S. Udawatte, Kang S. Wook, Yue Wang, Silvia Manzanero, Thiruma V. Arumugam, Chaminda J. Seneviratne

**Affiliations:** National Dental Research Institute Singapore (NDRIS), National Dental Centre Singapore, Oral Health ACP, Duke-NUS, Singapore; Department of Physiology, Yong Loo Lin School of Medicine, National University of Singapore, Singapore; Institute of Molecular and Cell Biology, A*STAR, Singapore; Jamieson Trauma Institute, Royal Brisbane and Women’s Hospital, Metro North Hospital and Health Service, Brisbane, Queensland, Australia; Department of Physiology, Anatomy & Microbiology, School of Life Sciences, La Trobe University, Australia; Oral Health ACP, Duke-NUS Medical School, Singapore

**Keywords:** gut microbiome, antifungal, 16S rRNA, ITS2, SM21, mycobiome, bacterium

## Abstract

The effects of antifungal agents on the human microbiome can be challenging to study due to confounding factors such as the underlying disease states and concomitant use of antibiotics and other therapies. We elucidated longitudinal modification of gut microbiome in response to a short course (5 days) of antifungal treatment in healthy male Sprague-Dawley (SD) rats by sequencing 16S rRNA V1–V3 and ITS2 hypervariable regions. SD rats were randomized into a control group and three antifungal treated (AT) groups including Amphotericin B (AmB), voriconazole and, our novel antifungal drug candidate SM21 once per day for 5 consecutive days. Fecal samples were collected at three different time points (day 0, day 1 and day 5). Microbial communities of both bacteria and fungi were compared between conditions. In silico analysis of differential microbial abundance and the predictive functional domains of microbial communities was further done by inferring metabarcoding profiles from 16S data. AT animals exhibited significant change in bacteriome alphadiversity although no divergence in community structure (beta-diversity) was observed compared with respective control groups (day 0). Specific bacterial clades and taxa were longitudinally and significantly modified in the AT animals. The AT bacterium of AmB and SM21 was particularly enriched in probiotic *Lactobacillus* strains including *L. reuteri*. The key pathways overrepresented in the bacteriome under AT animals were linked to cellular processes, environment information processing and metabolism. Moreover, AT treated mycobiome diversity decreased longitudinally with insignificant variations along the time course; different fungal taxa dominating at different timepoints in a wave-like fashion. However, acute antifungal treatments could not alter healthy gut microbial community structure. Hence, the healthy gut microbiome is capable of resisting a major dysbiotic shift during a short course of antifungal treatment.

## 1. Introduction

The gut microbiome plays an essential role in human health and disease. Various factors such as age, sex, dietary composition, genetic variations and pathological states influence the composition and abundance of the gut microbiome (Hopkins *et al.* 2001; Turnbaugh *et al.* 2010; Shetty, Marathe, and Shouche 2013; Goodrich *et al.* 2014). The gut microbiome aids the host to orchestrate the immune responses and to protect against invading pathogens (Holmes *et al.* 2011). Antibiotic treatments have been shown to modulate the structure and functionality of the gut microbiome (Ferrer *et al.* 2017). Previous studies have established that broad-spectrum anaerobic antibiotics such as ceftriaxone and ticarcillin-clavulanic acid are associated with increased yeast composition in the gut microbiome compared to antibiotics with poor anaerobic activity (e.g. ceftazidime, aztreonam, and imipenem-cilastatin) (Goodman, 1993). On the other hand, depletion of commensal intestinal fungi through the chronic use of antifungals has been shown to trigger the growth of pathogenic bacterial microbiota, which in turn exacerbates disease conditions in rodent models (Qiu *et al.* 2015).

Fungal infections commonly require the use of systemic antifungal agents. For instance, candidemia, which is a leading nosocomial infection (Strollo *et al.* 2017), requires amphotericin B-based preparations, azole and echinocandin antifungals for a considerable period of time. Echinocandins, voriconazole and Amphotericin B (AmB) (deoxycholate and lipid formulations) are commonly used as systemic antifungal agents (Keane *et al.* 2018). We have recently discovered the new antifungal small molecule SM21 which has shown promising effects against systemic candidiasis *in vivo* (Wong *et al.* 2014). Despite wide clinical use of antifungal agents, studies on their effect on the healthy gut microbiome are limited. To address this research gap, we investigated the longitudinal modification of gut microbiome in response to a short course of the commonly used antifungals AmB and voriconazole, and the novel antifungal agent SM21 in healthy Sprague-Dawley rats.

## 2. Methodology

### 2.1 Animals and experimental design

Thirty-six healthy five-week-old male Sprague-Dawley (SD) rats weighing 300–400g were purchased from Charles River Laboratories (Wilmington, MA, USA) (Strain Code # 400). The rats were fed a commercial pellet diet and deionized water ad libitum and housed two per cage at 20 °C ±2 °C and 50 to 70% relative humidity with a 12:12 hour light-dark cycle. After 2 weeks of acclimatization, the rats were weighed and randomized into three experimental and control groups and each group housed in separate cages for sampling of fecal pellets to avoid coprophagy-induced cross-contamination of samples, with three male rats per each condition (Figure 1). The groups were: AmB (Sigma-Aldrich, Cat # A9528, St. Louis, MO, USA), SM21 (PubChem CID: 5351098- [4-[2-(2,6-ditert-butylpyran-4-ylidene) ethylidene]cyclohexa-2,5-dien-1-ylidene]-dimethylazanium;perchlorate],

**Figure 1:**
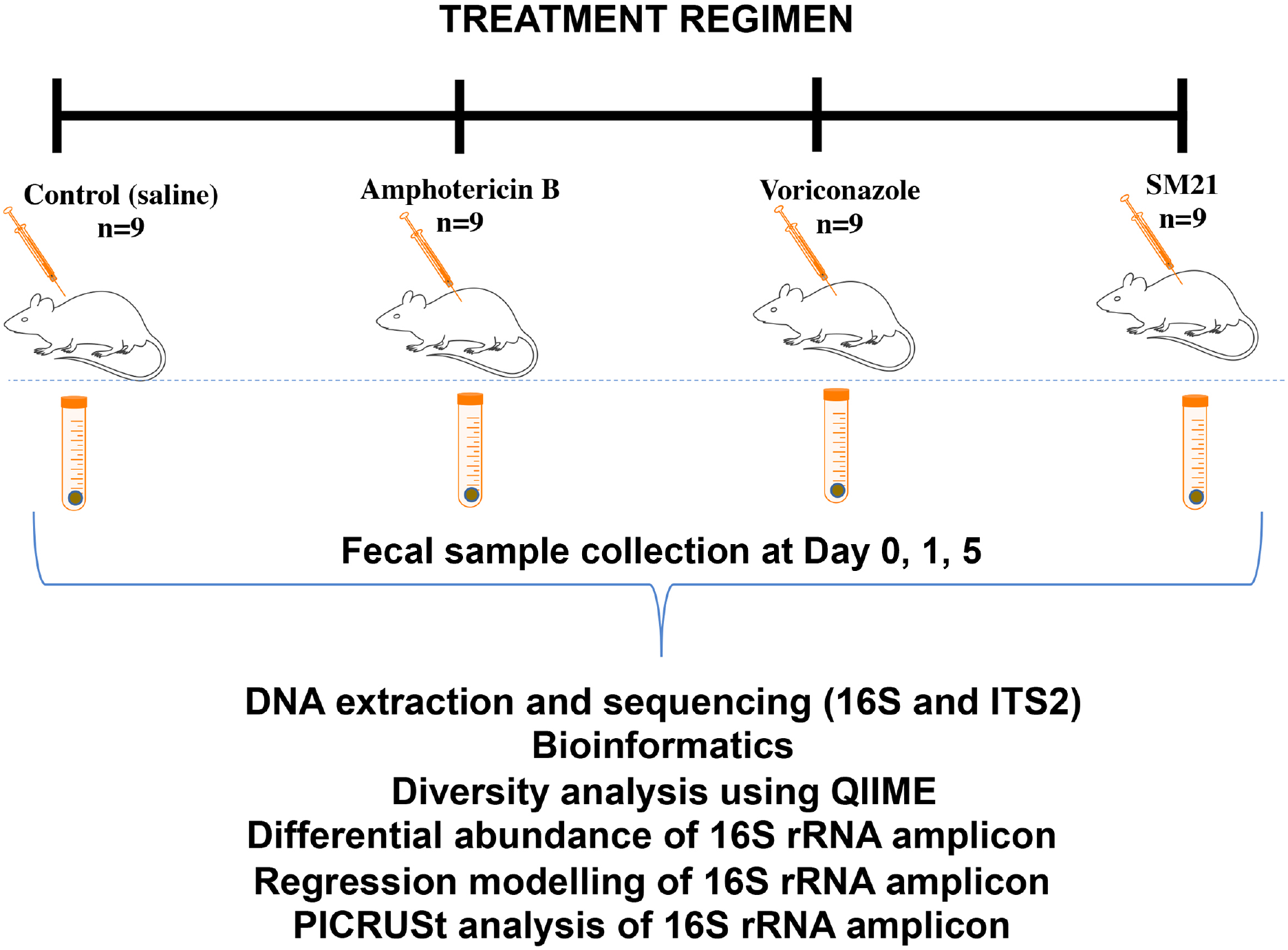
Experimental summary. Diagram of Antifungal dosing and fecal sample collection schedule. A cohort of 9 rats per group were treated once per day for five consecutive days, with sample collection at three different time points (n=3 per time point), followed by downstream analysis process.

ChemBridge, San Diego, CA, USA), voriconazole (Sigma-Aldrich, Cat # PZ0005, St. Louis, MO, USA), and vehicle control. Sterilized antifungals (1.5 mg per kg body weight) were administered once per day by intravenous injection, for 5 consecutive days. The control group received an equal volume of sterilized 0.9% saline). All animal-related procedures were conducted in an Association for Assessment and Accreditation of Laboratory Animal Care (AAALAC), an accredited facility at the National University of Singapore, and approved by the Institutional Animal Care and Use Committee (Approval number: R17-0327).

### 2.2 Sample collection

Fecal samples were collected from all rats (n=3 per each condition) at day 0 (before treatment), day 1 and, day 5 post antifungal treatment along with untreated controls. The rats were sacrificed on day 5 by administration of CO_2_ and cervical dislocation. Samples were placed in sterile centrifuge tubes immediately after collection and sealed to avoid cross-contamination between samples. All fresh fecal samples were stored in liquid nitrogen and then transported to the laboratory in a mobile refrigerator and frozen at −80°C until the extraction of DNA.

### 2.3 DNA extraction and sequencing

Total bacterial and fungal DNA was extracted using the QIAamp FastDNA Stool Mini Kit (Qiagen, Germany) according to the manufacturer’s protocol. DNA integrity was evaluated by gel electrophoresis (Wang *et al.* 2012). The V3-V4 region of the bacterial 16S rRNA gene was amplified by PCR using the primers 515F (5’ - CCTAYGGGRBGCASCAG-3’) and 806R (5’-GGACTACNNGGGTATCTAAT-3’) and the internal transcribed spacer 2 (ITS2) region of the fungal genome was amplified by PCR using the primers F (5’-GCATCGATGAAGAACGCAGC-3’) and R (5’-TCCTCCGCTTATTGATATGC-3’) (Bengtsson-Palme *et al.* 2013). Indexed adapters were added to the ends of the primers. All PCR reactions were carried out with Phusion® High-Fidelity PCR Master Mix (New England Biolabs, UK). The mixture of PCR products was purified using Qiagen Gel Extraction Kit (Qiagen, Germany). Sequencing libraries were generated using NEBNext® UltraTM DNA Library Prep Kit for Illumina, following manufacturer’s recommendations, and index codes were added. The library quality was assessed on the Qubit® 2.0 Fluorometer (Thermo Scientific, Singapore) and Agilent 2100 Bioanalyzer system (Agilent, Singapore). Finally, the library was sequenced on an Illumina platform and 250 bp paired-end reads were generated.

### 2.4 Microbiome sequence data processing

The overlapping regions between the paired-end reads were merged using FLASH (Magoč and Salzberg, 2011) and raw reads were quality filtered under specific filtering conditions to obtain high-quality clean tags on the basis of the Quantitative Insights into Microbial Ecology (QIIME, Version 1.7.0) quality control process (Caporaso *et al.* 2010). Sequences that were less than 200 bp in length or that contained homopolymers longer than 8 bp were removed. The chimera sequences were detected by comparing tags with the reference database RDP using the UCHIME (Edgar *et al.* 2011) algorithm and then removed (Haas *et al.* 2011). The effective sequences were then used in the final analysis. Sequences were grouped into operational taxonomic units (OTUs) using Uparse software (Uparse v7.0.1001). Sequences with ≥97% similarity were assigned to the same OTUs. The representative sequence for each OTU was screened for further annotation (Edgar, 2013). They were then taxonomically classified to different levels (phylum, class, order, family, genus, and species) using the Ribosomal Database Program (RDP) classifier (http://greengenes.lbl.gov/cgi-bin/nph-index.cgi) (Wang *et al.* 2007). OTU abundance data were normalized using a standard of sequence number corresponding to the sample with the least sequences. The output normalized data was used as the basis for analysis of alpha diversity and beta diversity.

### 2.5 Statistical analysis

Samples were rarefied to 75894 (read count of sample with lowest non-chimeric read) sequences. The rarefaction curve was plotted on the observed species using a QIIME script (alpha_rarefaction.py) and an in-house R script (unpublished). Alpha diversity is used to analyse complexity of biodiversity for a sample through indices - including Observed-species, Chao1 and Shannon. Good-coverage rarefied samples using for richness and diversity indices of the bacterial and fungal communities and displayed with R software (Version 2.15.3). Alpha diversity indices are presented as means ± SD. The differences in Alpha diversity indices and relative abundances between groups of the top 10 phyla and genera were calculated using the independent sample t-test (for normally distributed data) or the Mann-Whitney U-test (for non-normally distributed data). Beta diversity was calculated using unweighted UniFrac and nonmetric multidimensional scaling (NMDS), after which intra-group and inter-group beta distance boxplot diagrams were generated. A one-way analysis of similarity (ANOSIM) was performed to determine the differences in bacterial and fungal communities among groups (Clarke and Gorley, 2006). Unweighted Pair-group Method with Arithmetic Means (UPGMA) clustering was performed as a hierarchical clustering method to interpret the distance matrix using average linkage and was conducted by QIIME. A P-value < 0.05 was considered statistically significant, and P-value < 0.001 was considered extremely significant.

### 2.6 Differential analysis of 16S rRNA gene using LEfSe

A linear discriminant analysis coupled with the effect size (LEfSe) algorithm was used to identify the bacterial OTUs that longitudinally and significantly differed among the four groups (untreated control, AmB, voriconazole and SM21) based on the OTU relative abundance values using Galaxy workflow framework 2. Briefly, the algorithm first used the non-parametric factorial Kruskal-Wallis (KW) sum-rank test to detect the taxa with significantly different abundances, followed by pairwise Wilcoxon tests to detect biological consistency between the two groups. Finally, a linear discriminant analysis (LDA) score was used to estimate the effect size of each differentially abundant feature. A size-effect threshold of 4.0 on the logarithmic LDA score was used for discriminative functional biomarkers.

### 2.7 Regression modelling of 16S rRNA gene using Random Forest (RF)

To re-validate in identifying the fecal bacterial OTUs that differentiate among four groups (antifungal treatment and untreated control) by end of treatment day 5, the machine-based learning algorithm Random Forests was used in R v3.2.5, Random forest package v4.6.x. The most differently abundant genera derived from LEfSe used to generate the Random Forests-generated model discriminating all three antifungal treatment and untreated control. 10-fold cross-validation was performed to search to select the optimal subset of bacteria discriminating among all four groups. The final model accuracy is taken as the mean from the number of repeats. The classification performance of the final model was assessed with OOB error rate. The “gplots” was used to plot OTU relative abundances as heat maps. The importance of a variable was calculated as the mean decrease in accuracy.

### 2.8 Predictive metagenomic analysis of 16S rRNA gene using PICRUSt

The functional capacity of the gut microbiome was estimated by inferring metabolic functionality from the 16S rRNA gene sequencing data using the Phylogenetic Investigation of Communities by Reconstruction of Unobserved States (PICRUSt) v1.0 software. Z-scores were calculated to construct a heatmap of the relative abundance of the pathways in each group with the formula z = (x - μ)/σ, where x is the relative abundance of the pathways in each group, μ is the mean value of the relative abundances of the pathways in all groups, and σ is the standard deviation of the relative abundances. Phylogenetic Investigation of Communities by Reconstruction of Unobserved States (PICRUSt) analysis was performed to identify Kyoto Encyclopedia of Genes and Genomes (KEGG) metabolic pathways potentially affected by groups of bacteria based on taxonomy obtained from the Greengenes reference database1 (DeSantis *et al.* 2006) using Galaxy workflow framework 2. Pathway topology analysis was conducted by using the pathway data in *Rattus norvegicus* pathway libraries (including 81 metabolic pathways). Differential abundance analysis utilized the nonparametric permutation difference test for a diversity and PICRUSt values or the negative binomial test (Kanehisa *et al.* 2014) for taxonomic count data.

## 3. Results

### 3.1 Sequence analysis

A total of 36 samples were collected, comprising three animals per each condition, four groups (AmB, SM21, voriconazole and untreated control) and three time points (day 0, day 1 and day 5). For the 16S rDNA gene, a total of 120,301—150,685 raw paired-end sequence numbers (mean 147,617 ± SD 7511.66) and analyzed sequences (mean length = 410 bp) were obtained from each sample. For ITS2, a total of 176,897—288,978 raw paired-end sequence numbers (mean 195,663 ± SD 4350.81), and analyzed sequences (mean length = 425 bp) were acquired from each sample. A total of 1372 operational taxonomic units (OTUs) and 1555 OTUs were obtained from 16S rDNA and ITS2 regions, respectively, at a 97% sequence-similarity level. To evaluate the species coverage identified by our sequencing effort, rarefaction analysis was performed (Supplemental Figure 1). Although the rarefaction curves from both 16S rDNA and ITS2 did not achieve the even stage, their terminal slopes were rather low, suggesting that the sequencing detected the majority of the species.

### 3.2 The dynamics of gut microbiome abundance and distribution

The bacterial distribution of untreated control rats was dominated by Firmicutes (70-75%) and Bacteroidetes (20-25%) followed by Proteobacteria (1-2%) and Verrucomicrobia (0.5-1%) (Figure 2A). *Romboustia, Parabacteriodes, Ruminoccuss*, and *Lactobacillus* were observed to be the dominant genera (Figure 2B). When assessing the effect of specific antifungals, AmB caused a significant temporal shift (p < 0.05) in the ratio of Firmicutes to Bacteroidetes across the treatment time points (D.1-D.5). Further, rats treated with either AmB or voriconazole showed a significant (p < 0.05) relative decrease in *Romboutsia* and *Ruminoccuss* populations. SM21 treatment did not significantly affect bacterial taxa across the treatment time points.

**Figure 2:**
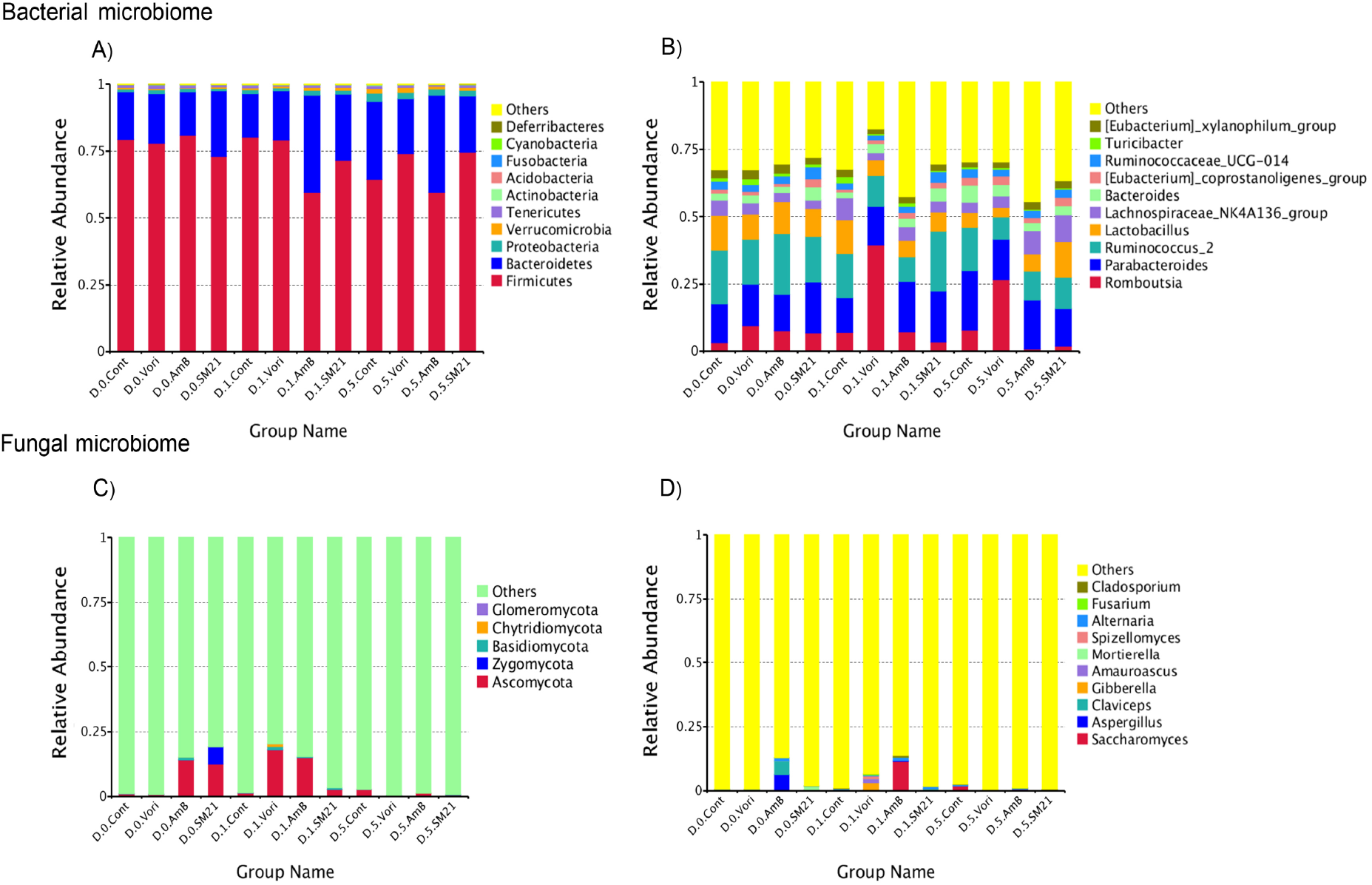
Histograms depicting the dynamics of gut microbiome abundance and distribution of the top 10 taxanomic level. The x-axis represents treatment groups and the y-axis represents relative abundance presented as a fraction of one. Relative abundance of the top 10 phyla, (A) bacterial microbiome and (C) fungal microbiome. Relative abundance of the top 10 genera, (B) bacterial microbiome and (D) fungal microbiome. Only the top 10 taxa in the abundance are shown in the figure; other taxa were combined as “Others.”

Distribution of gut mycobiome in untreated control rats was dominated by Ascomycota (5.9%), Zygomycota (0.6%) and Basidomycota (0.3%), followed by Chytridiomycota (0.1%) and Glomeromycota (0.1%) (Figure 2C); *Saccharomyces, Aspergillus, Claviceps* and, *Gibberella* were the dominant genera (Figure 2D) along with a substantial number of unknown abundant lineages. It should be noted that a significant proportion of reads remained unclassified at the each taxonomical level. These observations indicate the presence of several hitherto unknown fungal taxonomic groups in the rat gut and highlight the limitations of the existing fungal sequence databases.

### 3.3 Core gut microbiome changes

Heat maps (Figure 3A) were constructed to evaluate the effect of core bacterial OTU abundance under each antifungal treatment. Comparing D.1 and D.5, we observed that antifungal treatment enriched the abundance of bacterial OTUs. To examine the change of common core microbiome longitudinally, we defined a core as the group of members common to each day and represented the core by overlapping areas in a Venn diagram, at 97% identity. As shown in Figure 3B, the number of OTUs shared among the four groups increased through the time course with 334 on D.0, 368 on D.1 and 381 on D.5. Instead, the number of core fungal OTUs decreased across the treatment time points under antifungal exposure from 92 on D.0 to 83 on D.1 and 72 on D.5 (Figure 3D). Mycobiome comparison of samples collected one day after antifungal treatment revealed a relative increase in abundance (>0.1%) in a few clades (Ex: *Saccharomycestes, Giberella, Fusarium, Alternaria etc*.) which were not dominant members of the communities in untreated controls or prior to antifungal treatment (D.0) (Figure 3C & E). However, these dominant lineages decreased sharply in abundance in samples collected at day 5 (Figure 3C). Overall, taxonomies account for less than 0.3% of total core gut fungal mycobiota in SD rats.

**Figure 3:**
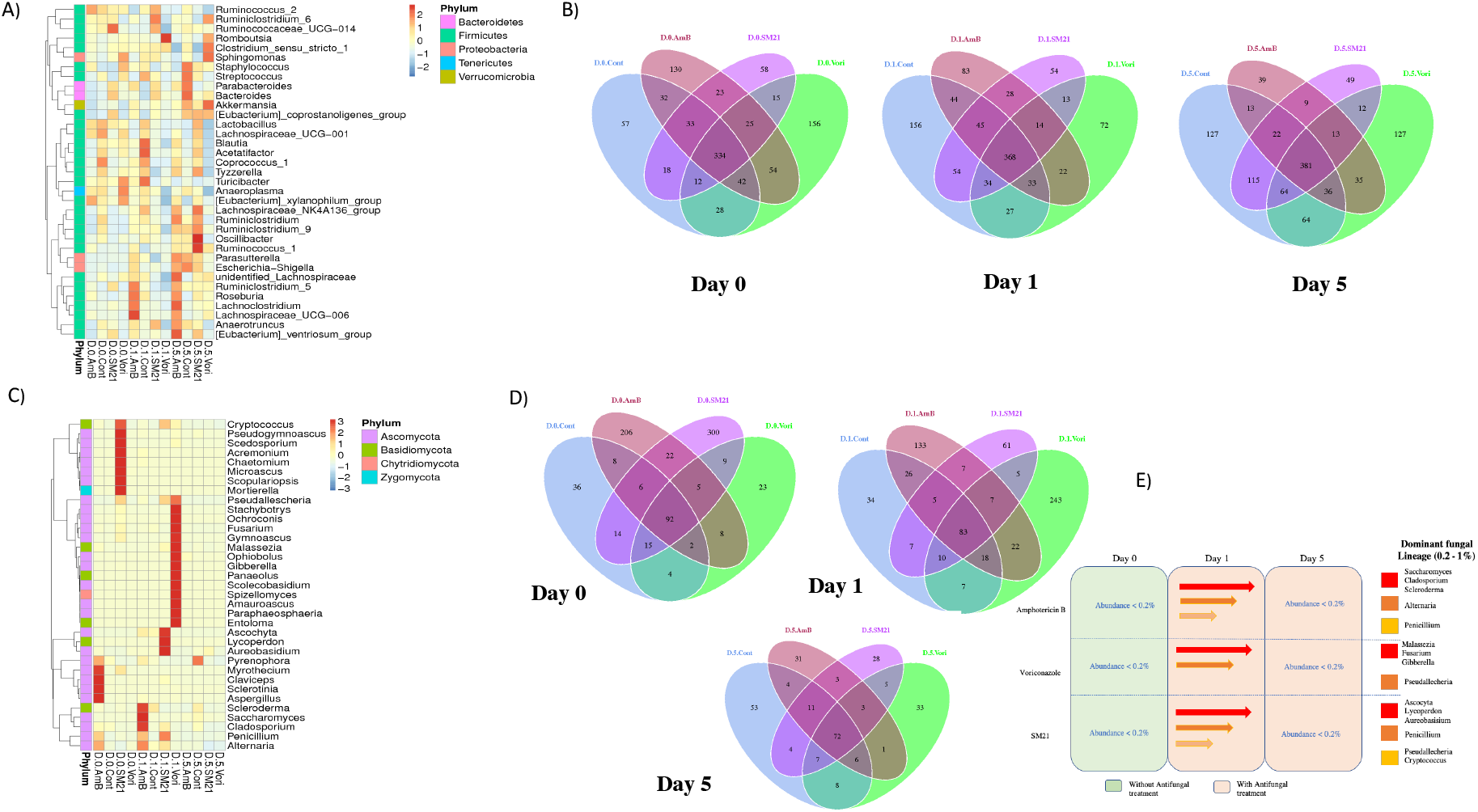
Core gut microbiome changes depicted by OTU heatmap and Venn diagram in samples treated with voriconazole, Amphotericin B, and SM21 at each treatment time point (day 0, 1, and 5). Heatmap depicting abundance of the top 35 OTUs by overall abundance across samples. Heatmap was constructed using normalized log10 abundance of each OTU in each sample. To the right of the heatmap is a list of OTUs generated by RDP database assignment or best-hit classification where appropriate. The higher the relative abundance of an OTU in a sample, the more intense the color at the corresponding position in the heatmap. Heatmap by (A) bacterial microbiome and (C) fungal microbiome. The Venn diagrams show the numbers of OTUs (97% sequence identity) that were shared or not shared by all different samples. Venn diagrams by (B) bacterial microbiome and (D) fungal microbiome. (E) Longitudinal variation in fungal abundance in antifungal treatment. Rows show treatment groups, columns show time points (D.0, D.1 and D.5) and dominant fungal lineages are shown on the right.

### 3.4 Microbial diversity changes

Next, we evaluated whether the longitudinal changes of microbiome lineages reported above were paralleled by the changes in microbial diversity associated with each antifungal treatment compared to respective controls. Estimated and observed species diversity (Shannon) and richness (Chao 1) were determined. In the bacterial community, Shannon’s index, which accounts for both the number of OTUs and their relative abundance, did not differ significantly between groups in untreated control samples throughout the time course. However, group-specific changes across the treatment time points indicated diversified OTU abundance. The Shannon diversity index was higher with significant difference in AmB-treated group between day 0-1, (P = 0.0242) day 0-5 (P= 0.0215); SM21 treatment between day 0-5 (P = 0.038) and, voriconazole between day 0-1 (P = 0.011) (Figure4A). However, the Chao1 index remained similar amongst treatment groups with respect to day 0 groups, revealing no differences in the expected species richness of the bacterial microbiome in each treatment group (Fig. 4B). The microbial diversity and richness of fungal communities under acute antifungal exposure remained unchanged, as showed by Shannon and Chao I indices (Fig. 4C & D) Good’s coverage estimator for each group was over 95%, indicating that the current sequencing depth was sufficient to saturate both bacterial and fungal diversity (Supplementary Table. 4 & 5).

**Figure 4:**
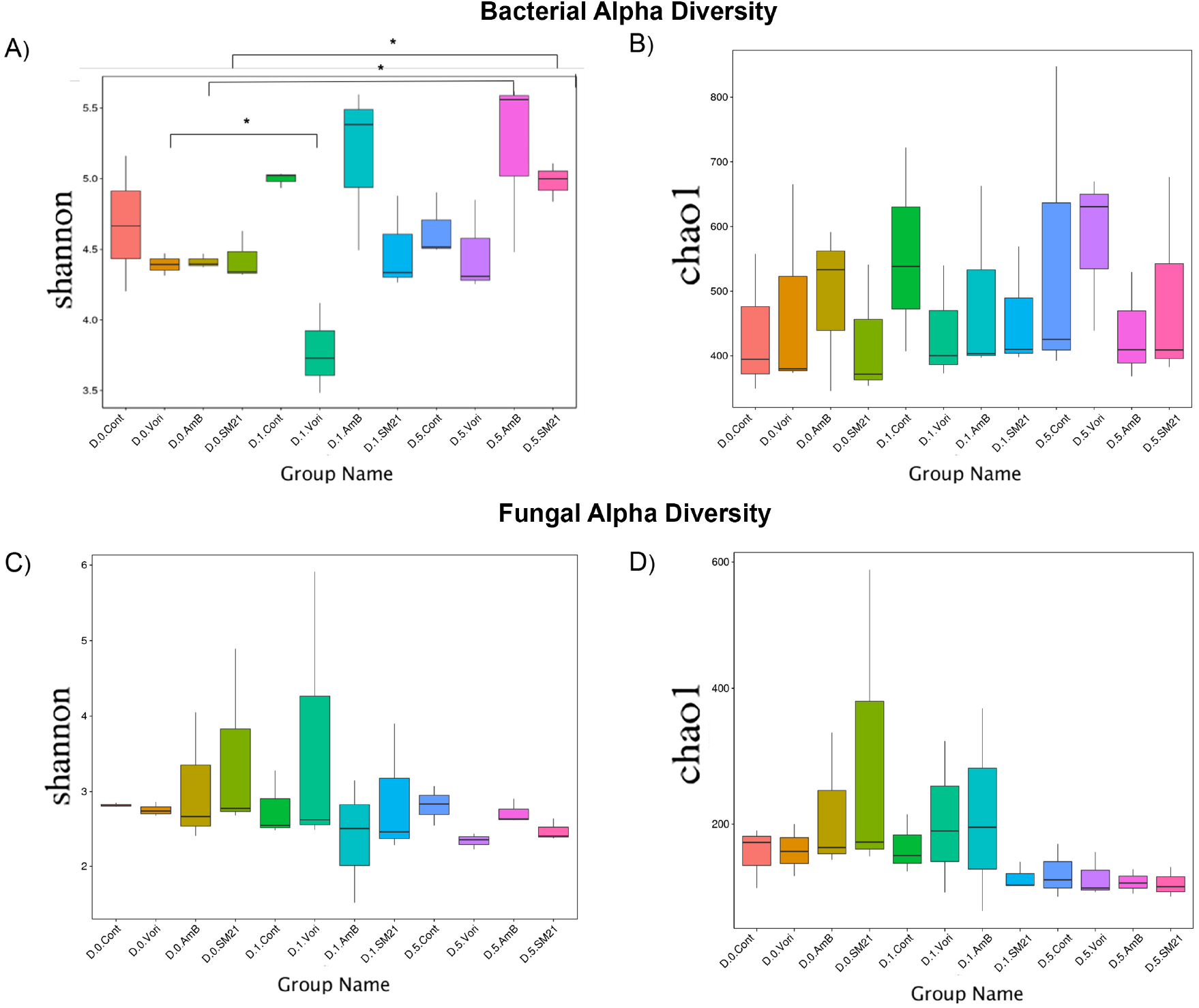
Boxplot of Alpha-diversity indices (Shannon and Chao 1). Alpha diversity indices are composite indices reflecting abundance and consistency. Shannon and Chao I indices of; (A) & (B) bacterial microbiome, and (C) & (D) fungal microbiome. The greater the Shannon index, the higher the diversity of the microbiota; The greater the Chao I index, the higher the expected species richness of the microbiome. Boxes represent the interquartile range (IQR, 25th and 75th percentiles), and the horizontal line inside the box defines the median. Whiskers represent the lowest and highest values within 1.5 times the IQR from the first and third quartiles, respectively. “*” P < 0.05 (t-test, wilcox and Tukey tests).

### 3.5 Microbial community structure changes

We next evaluated the longitudinal differences of microbial community structure between treatment groups and untreated controls using Non-Metric Multi-Dimensional Scaling (NMDS). We employed an ANOSIM analysis on weighted UniFrac distance to evaluate whether variation among treatment groups is significantly larger than variation within treatment groups.

Significant treatment-associated clustering was not observed in both bacterial and fungal community structures depicted by NMDS plots (Figure 5A; Supplemental Table 6). However, the inference of the phylogenetic tree based on UPGMA (Unweighted Pair Group Method with Arithmetic Mean) hierarchical clustering analysis has shown the diversified pattern of bacterial Phyla, showing day 5 treatment and respective day 0 control groups of voriconazole and AmB located more towards on the terminal branch ends of the tree (Figure 5B). Further, ANOSIM similarities matrices (R > 0, P_anosim_ > 0.05) shows that inter-group differences are greater than intra-group differences (Supplemental Figure 2). Of these, the bacterial species showing significant inter-group difference observed to be between day 0 and day 5 under AmB and voriconazole treatment. The significant taxon defined by the shared presence of both groups belonging to the genus *Romboustia* (P= 0.028 and P= 0.040; respectively).

**Figure 5:**
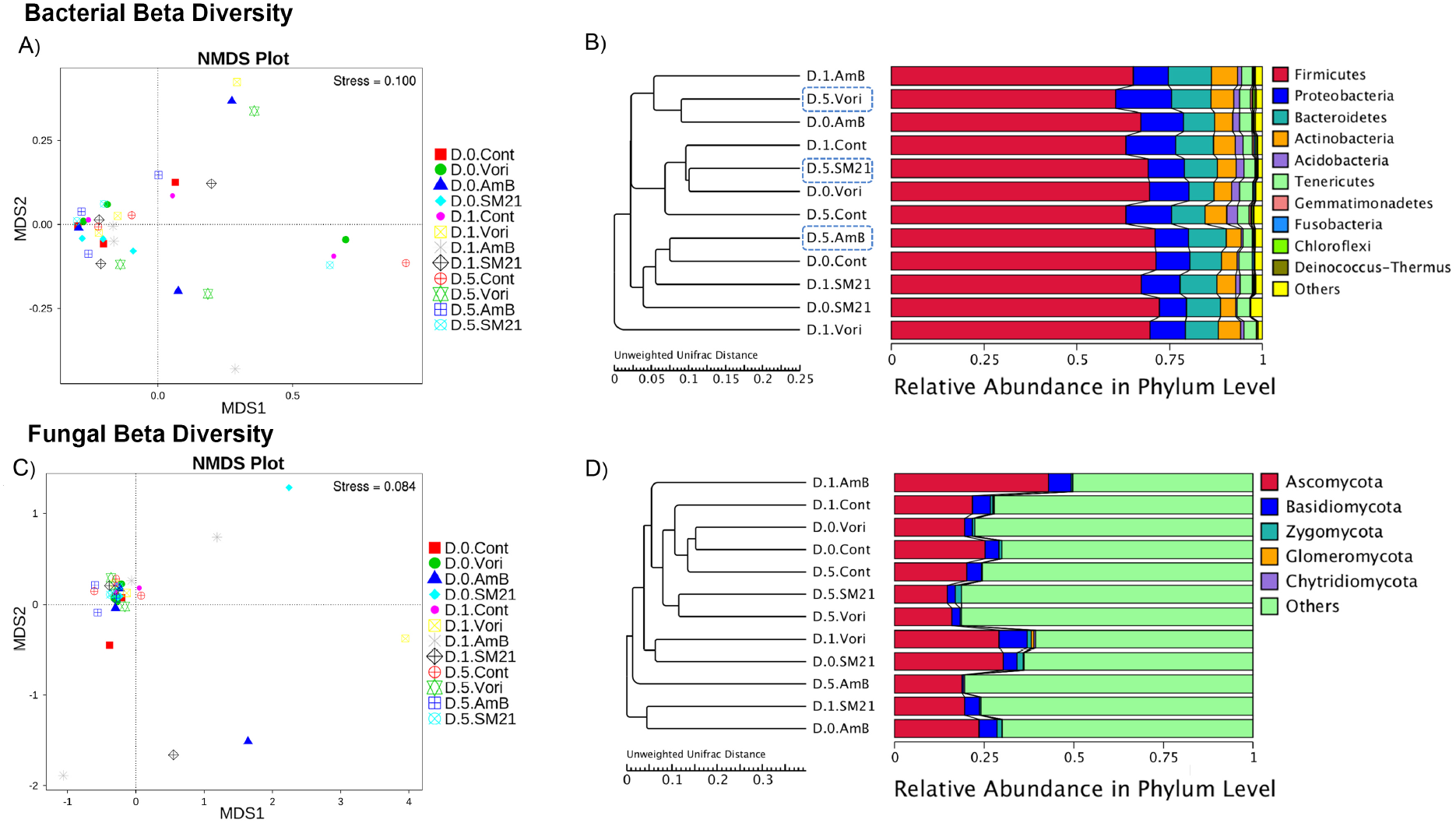
Non-metric multidimensional scaling (NMDS) and UPGMA (Unweighted Pair Group Method with Arithmetic Mean) analysis of beta diversity. Each point in the NMDS graphs represents one sample, and different colours represent different groups. The distance between points represents the level of difference. Stress lower than 0.2 indicates that the NMDS analysis is reliable. (A) Bacterial beta diversity, (B) fungal beta diversity. UPGMA tree of unweighted UniFrac distances; C) bacterial diversity and D) fungal diversity. They were displayed with the integration of clustering results and the relative abundance of each sample by phylum level. The diversified pattern of bacterial Phyla, showing all day 5 treatment groups and corresponding day 0 groups of voriconazole and AmB located on the terminal branch ends of the tree on figure B (blue dashed line squares on all day 5 groups).

Taken together, the results show that acute antifungal treatments did not alter healthy gut microbial community structure, but did affect specific bacterial taxa. These changes in bacterial taxa under each antifungal treatment have been considered for further association of differential bacterial abundance analysis and predictive metabolic functions.

### 3.6 Differential bacterial abundance

Utilizing the LEfSe algorithm, we identified total 17 different taxa that differed in abundance for all three antifungal treatments by end of day 5 (Supplemental table 9). Of these, five taxa had significantly higher abundances in both AmB and voriconazole treatment and two taxa had higher abundances in SM21 treatment (Figure 5B). The phylum Firmicutes and genus *Romboutsia* were particularly abundant in voriconazole treatment (LDA Score > 4.8). Moreover, LEfSe highlighted the overabundant Firmicutes within the Clostridia clade (family *Peptostreptococcaceae*) under voriconazole treatment (Figure 6A & B). At the species level, *Lactobacillus reuteri* was significantly enriched in SM21 treatment (LDA Score > 4.8) and at the genus level, *Lactobacillus* and *Lachnospiraceae_NK4A136_group* (LDA Score > 4.8) were prominently abundant in AmB treatment (Figure 6B & C).

**Figure 6:**
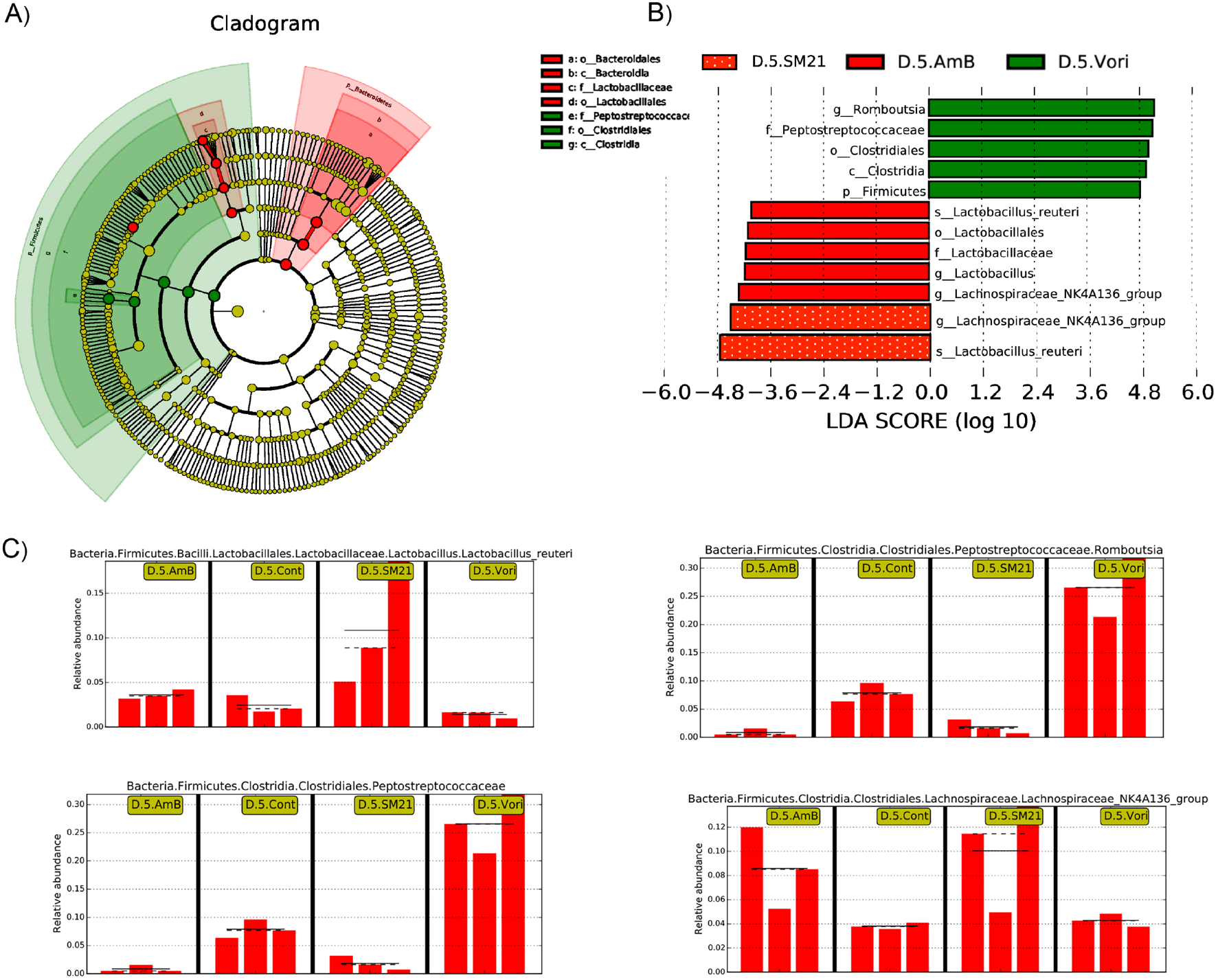
Differential gut bacterial abundance using LEfSe analysis. (A) Cladogram representation of the differentially abundant taxa, in each treatment group with respect to the control groups. The root of the cladogram represents the domain bacteria. The size of each node represents their relative abundance. No significantly different taxa are labelled in yellow. Significantly different taxa are labelled with the colour of main two taxa differed in the treatment. (B) The top 10 taxa with the largest effect sizes (LDA Score > 4.5) are presented in the figure. The length of the histogram represents the LDA score; i.e., the degree of influence of species with significant difference between different groups after 5 day of post treatment of each antifungal. (C) The x-axis represents antifungal groups and the y-axis represents relative abundance of most significant species between the 3 treatment groups and untreated control (p < 0.05).

### 3.7 Predictive model for antifungal treatment on gut bacteriome signature

Next we validated the above findings using regression-based Random Forest (RF) to identify to which extent the observed significantly abundant bacteria correlate between antifungal exposure and untreated controls by end of the treatment (day 5). A random forest algorithm was used to re-assess the most predictive 17 taxa based on the LEfSe as described in methodology. Two highly discriminating genera with the highest calculated variable importance measurement were selected as the most significant taxa segregating between this two cohorts (Supplemental figure 4). Among the genera identified by the Random Forest classifier as contributing the most to the group classification model were *Lactobacillus reuteri and Romboutsia*. Using LEfSe and RF, *Romboutsia* and *Lactobacillus reuteri* are consistently identified as showing the most important differential genera; marked by higher mean decrease in accuracy.

### 3.8 Functional prediction of gut bacteriome in reponse to antifungal treatment

To understand the functional role of the gut microbiome following antifungal treatment, the normalized PICRUSt gene family counts were plotted as heatmaps. The plots revealed several hot spots for each treatment group. Five KEGG pathways, namely Transport and Catabolism, Glycan Biosynthesis and Metabolism, Metabolism of Cofactors and Vitamins, Energy Metabolism, and Biosynthesis of Other Secondary Metabolites, were overrepresented in AmB post treatment samples. Amino Acid Metabolism and Signal Transduction were exclusively overrepresented in SM21 treatment on day 1 and day 5, respectively. On the other hand, ten KEGG pathways in the voriconazole group were overrepresented on day 1 including Signaling Molecules and Interaction, Cellular Processes and Signaling, Carbohydrate Metabolism, Xenobiotic Biodegradation and Metabolism. The representation of these pathways decreased on day 5 (Figure 7A & B). The key pathways overrepresented in the gut bacteriome under antifungal treatment were linked to cellular processes, environment information processing and metabolism (Figure 7C). In contrast to the NMDS plot based on bacterial taxonomic abundance (Figure 5), the PCA plot based on functional pathways revealed distinct clustering of treatment samples over control groups (except day 1 post-treatment with SM21) (Figure 7D).

**Figure 7:**
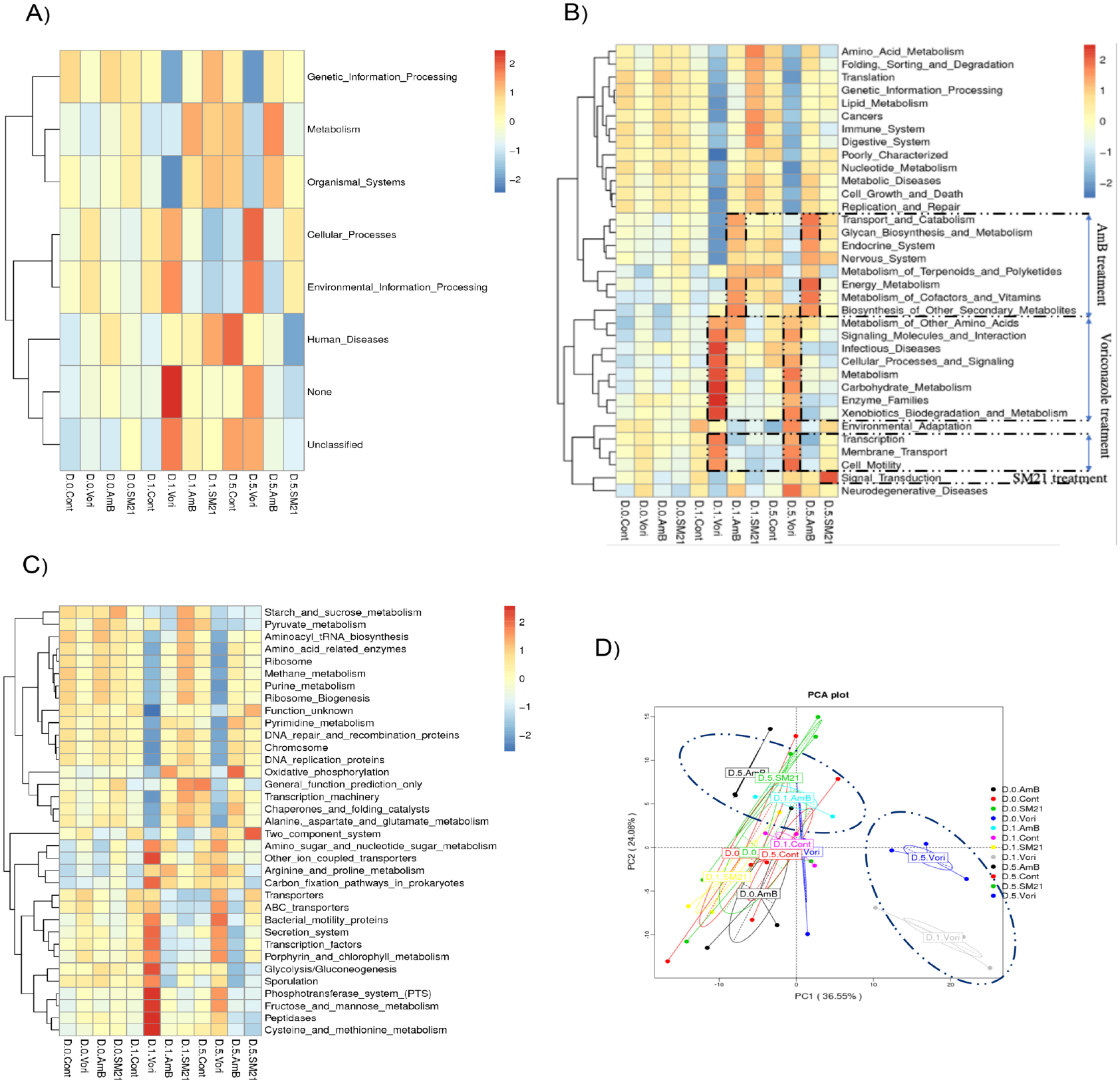
Functional prediction of gut bacteriome changes using PICRUSt. A) KEGG Functional Hierarchy level 1, B) KEGG Functional Hierarchy level 2 and, C) KEGG Functional Hierarchy level 3. Predicted function of the gut bacterial microbiota in AmB, voriconazole, SM21 treatment and control groups. Z-scores were calculated to construct a heatmap of the relative abundance of the pathways in each group. The columns represent groups at each treatment time point, and the rows depict metabolic pathways. The color coding is based on differentially expressed genes leading to changes in abundance of pathways. D) PCA plot describing functional inferences (PICRUSt) of bacterial communities across antifungal treatment groups and time points. The colors indicate the treatment time points of the same antifungal treatment in a given site. Ovals show two distinct clusters of samples, one of samples treated with voriconazole and the other of samples treated with AmB and SM21 (day 5 only).

## 4. Discussion

In this study, the healthy gut microbiome was not significantly influenced by acute antifungal treatment in the SD rat model, although specific bacterial taxa showed significant differences with AmB and voriconazole treatment. The modulation of probiotic *Lactobacillus* strains including *L. reuteri* in response to AmB and antifungal drug candidate SM21 treatment could be a response aimed at restoring a healthy gut microbiome. Instead, gut mycobiome diversity decreased longitudinally with antifungal treatments with insignificant variation along the time course, with different fungal taxa dominating at different timepoints in a wave-like fashion.

Despite changes in specific bacterial taxa, the overall healthy gut bacterial community structure was not affected by the antifungal drug treatment. Few studies have specifically investigated the effect of antifungals on the gut microbiome, and the data available focused mostly on disease-induced animal models (Qiu *et al.* 2016,2015; Botschuijver *et al.* 2017). However, our findings are similar to a study of mice treated with antifungals fluconazole and Amphotericin B, which also observed a stable commensal bacterial community structure with few changes of taxa even after several weeks of treatment (Wheeler *et al.* 2016). In our study, the gut bacterium prior to antifungal treatment was dominated by the phyla Firmicutes and Bacteroidetes. This remained stable during the course of voriconazole and SM21 treatment. However, AmB treatment significantly changed the bacteriome by day 5. In addition, we observed changes in bacterial diversity (Shannon index) with antifungal use, suggesting that there is a progressive depletion or colonisation of few bacterial taxa throughout the treatment. The gut bacterial genes of SD rats are known to have a higher pairwise overlap with humans (2.47%) than other animal models, including ninety-seven percent of the functional pathways in the human catalogue (Pan *et al.* 2018). Therefore, the findings of this study provide evidence base to investigate the effect of antifungals on gut microbiome in human clinical trials.

Among the taxa which increased in abundance after antifungal treatment, it is worth highlighting *Lactobacillus reuteri and*, *Lactobacillales are* well-known probiotic bacterium able to modulate the microbiome as well as the host’s immune system towards health (Thomas and Versalovic 2010; Yuying Liu et al; 2016; Llewellyn and Foey 2017). *L. reuteri* communicates with the host by inhibiting key signaling cascades such as NFkB-mediated pathways in the gastrointestinal tract. PICRUSt analysis revealed that SM21 treatment results in overrepresentation of KEGG pathways related to signal transduction. It is possible that *L. reuteri* which we identified as a taxonomic biomarker may be responsible for this functional change.

Furthermore, *L. reuteri* has been associated with other multiple metabolic pathways and has shown functional capacity to establish interactions between beneficial microbes and the human host (Saulnier *et al.* 2011). According to the foregoing studies, the major pathways associated with the abundance of *L. reuteri* alone or in combination with *Lactobacillales* are related to cofactors, vitamins, glycans, and signaling. In agreement with these studies, PICRUSt analysis showed an enrichment of pathways for the metabolism of cofactors and vitamins, and glycan biosynthesis and metabolism, following AmB treatment. Previous studies have shown that probiotic *Lactobacillus* species, in particular *L. reuteri* strains have a protective role against fungal infections (Guo *et al.* 2012; Matsubara *et al.* 2016; Seneviratne *et al.* 2016; Zangl *et al.* 2020). Therefore, our observations raise an intriguing possibility that the modulation of probiotic *Lactobacillus* strains, including *L. reuteri*, may be a response to repair the dysbiotic gut microbiome caused by AmB and SM21 treatment.

The genus *Romboutsia* in the gastrointestinal tract have been shown to encode a versatile array of metabolic capabilities for carbohydrate utilization, fermentation of single amino acids, anaerobic respiration and metabolic end products, based on comparative human genome analysis (Gerritsen *et al.* 2019). In the present study, PICRUSt pointed to pathways involved in the Metabolism of Carbohydrates and Amino acids (cystein and methionine) Peptidases, Fructose and Mannose, and Glycolysis could be linked to significant taxonomic biomarker of genus *Romboutsia* identified in voriconazole treatment. Further, RF predictions showed a highest correlation between *Romboutsia* modulation and voriconoazole treatment, followed by between modulation of *L. reuteri* and SM21 treatment. Although these correlations do not necessarily indicate interactions between each antifungal treatment and the metabolic potential of a single gut microbe, most of these metabolic pathways were not enriched in control samples during the experimental period, suggesting that gut dysbiosis is strongly associated with antifungal treatment. In addition, KEGG functional pathway variations and the PCA plot based on KEGG modules showed that most posttreatment microbiome changes are distinct. However, the structure and composition of intestinal microbiota with respect to functional activities are complex and multifactorial. Moreover, PICRUSt is only a tool for predicting functional genes; thus, further research is required to obtain more accurate gene function information by metagenomic analysis.

The foregoing data suggest that fungal and bacterial communities in the gut are co-dependent and that disruption of one affects the other. Fungal diversity composition changed radically over the time after one day of antifungal treatment, followed by a sharp decline at the end of treatment (day 5). However, fungal community structure did not change significantly by the short-course of antifungal treatment. The depletion of commensal fungal populations may require several weeks of exposure, as shown in a study using fluconazole and amphotericin B treatment in mice (Wheeler *et al.* 2016). Unexpectedly in our study, not just the different treatment groups, but also the control group, showed significant variation along the time course, with different fungal taxa dominating at different timepoints in a wave-like fashion. Similar observations have been made in mice where gut fungal mycobiome varied substantially over time receiving antibiotics as well as untreated control group (Dollive *et al.* 2013). Furthermore this study also showed that mice housed in different cages, but receiving the same treatment differed in their fungal microbiome. Hence, it was concluded that mice harbour individualised gut microbiota, irrespective of other factors such as diet or treatment groups (Dollive *et al.* 2013). This has also been observed in human studies. A human gut mycobiome study of 24 individuals at two sampling time points found that detection of the same fungus at both time points occurred less than 20% of the time (Hallen-Adams *et al.* 2015). Together with previous studies, our findings emphasize that the response of the gut mycobiome to antibiotics or antifungals varies substantially with treatment regimen, and other experimental factors. Our study also highlights that the data available on rat core mycobiome composition remains sparse. Our data in untreated control samples suggested gut commensal fungal proportions of SD rat in the range of 0.1–1.3%, with a median of 0.3%, dominated by Ascomycota, Zygomycota and Basidomycota and at genus level, dominated by *Saccharomyces*, *Aspergillus* and *Claviceps*. However, out of the 15 most abundant fungal genera found in a healthy human stool, we found only 6 (*Saccharomyces, Clavispora, Alternaria, Fusarium, Aspergillus, Penicillium*) in rat commensal core mycobiome in very low abundance, along with the absence of most abundant commensal genera of *Malassezia* and *Candida* in human.

Some limitations should be considered in the context of our investigation. First, due to the short treatment period, the community and diversity changes of the healthy gut microbiome under chronic exposure to antifungal treatment could not be determined. Second, we have not observed the restoration period of specific bacterial taxa to the original microbiota state (baseline) after withdrawal of antifungal treatment. Third, a significant proportion of uncharacterized fungal taxa were unavailable for examination in the database, which may impact the true estimation of fungal diversity in these samples.

In conclusion, the healthy gut microbiome is capable of resisting a major dysbiotic shift during a short course of antifungal treatment. However, there is a significant modulation of key few bacterial taxa in the gut microbiome. In particular, the modulation of probiotic *Lactobacillus* strains by AmB treatment and the induced growth of *Lactobacillus reuteri* by SM21 treatment warrant further investigations. Taken together, our results provide a new insight into the impact of a short course of antifungals on healthy gut microbiome, which will provide a basis for further investigation in clinical cohorts in future.

## Acknowledgement

We thank Novagene, Singapore for their role in assisting with data processing and analysis for publication.

## Author’s contributions

NU, SK, TA, and CS conceived and designed the study. NU, SK, and TA conducted experiments and data gathering. YW participated in the study design. NU designed, optimized, performed the sequencing experiments and analysis. NU, TA, SM and CS wrote the manuscript. Novagene performed the data processing. All authors critically reviewed and approved the manuscript.

## References

Botschuijver, Sara, Guus Roeselers, Evgeni Levin, Daisy M. Jonkers, Olaf Welting, Sigrid E.M. Heinsbroek, Heleen H. de Weerd, et al. 2017. “Intestinal Fungal Dysbiosis Is Associated With Visceral Hypersensitivity in Patients With Irritable Bowel Syndrome and Rats.” Gastroenterology 153 (4): 1026–39. https://doi.org/10.1053/j.gastro.2017.06.004.

Caporaso, J Gregory, Justin Kuczynski, Jesse Stombaugh, Kyle Bittinger, Frederic D Bushman, Elizabeth K Costello, Noah Fierer, et al. 2010. “Correspondence QIIME Allows Analysis of High-Throughput Community Sequencing Data Intensity Normalization Improves Color Calling in SOLiD Sequencing.” Nature Publishing Group 7 (5): 335–36. https://doi.org/10.1038/nmeth0510-335.

Clarke, K. R., and Gorley, R. N. 2006. Primer v6: User Manual/Tutorial. Plymouth: Plymouth Marine Laboratory.

DeSantis, T. Z., P. Hugenholtz, N. Larsen, M. Rojas, E. L. Brodie, K. Keller, T. Huber, D. Dalevi, P. Hu, and G. L. Andersen. 2006. “Greengenes, a Chimera-Checked 16S RRNA Gene Database and Workbench Compatible with ARB.” Applied and Environmental Microbiology 72 (7): 5069–72. https://doi.org/10.1128/AEM.03006-05.

Dollive, Serena, Ying Yu Chen, Stephanie Grunberg, Kyle Bittinger, Christian Hoffmann, Lee Vandivier, Christopher Cuff, James D. Lewis, Gary D. Wu, and Frederic D. Bushman. 2013. “Fungi of the Murine Gut: Episodic Variation and Proliferation during Antibiotic Treatment.” PloS One 8 (8). https://doi.org/10.1371/journal.pone.0071806.

Edgar, Robert C. 2013. “UPARSE: Highly Accurate OTU Sequences from Microbial Amplicon Reads.” Nature Methods 10 (10): 996–98. https://doi.org/10.1038/nmeth.2604.

Edgar, Robert C., Brian J. Haas, Jose C. Clemente, Christopher Quince, and Rob Knight. 2011. “UCHIME Improves Sensitivity and Speed of Chimera Detection.” Bioinformatics 27 (16): 2194–2200. https://doi.org/10.1093/bioinformatics/btr381.

Ferrer, Manuel, Celia Méndez-García, David Rojo, Coral Barbas, and Andrés Moya. 2017. “Antibiotic Use and Microbiome Function.” Biochemical Pharmacology 134 (September): 114–26. https://doi.org/10.1016/j.bcp.2016.09.007.

Gerritsen, Jacoline, Bastian Hornung, Jarmo Ritari, Lars Paulin, Ger T. Rijkers, Peter J. Schaap, Willem M. de Vos, and Hauke Smidt. 2019. “A Comparative and Functional Genomics Analysis of the Genus Romboutsia Provides Insight into Adaptation to an Intestinal Lifestyle.” BioRxiv, 845511. https://doi.org/10.1101/845511.

Goodman, Larry J. 1993. “Prospective Evaluation Oe Effects of Broad-Spectrum Antibiotics on Gastrointestinal Yeast Colinization of Humans.” Infectious Diseases in Clinical Practice 2 (3): 219–20. https://doi.org/10.1097/00019048-199305000-00018.

Goodrich, Julia K, Jillian L Waters, Angela C Poole, Jessica L Sutter, Omry Koren, Ran Blekhman, Michelle Beaumont, et al. 2014. “Article Human Genetics Shape the Gut Microbiome.” Cell 159 (4): 789–99. https://doi.org/10.1016/j.cell.2014.09.053.

Guo, Jiahui, B Brosnan, Ambrose Furey, E K Arendt, Padraigin Murphy, and Aidan Coffey. 2012. “Antifungal Activity of Lactobacillus against Microsporum Canis, Microsporum Gypseum and Epidermophyton Floccosum” 3 (2): 104–13.

Haas, Brian J., Dirk Gevers, Ashlee M. Earl, Mike Feldgarden, Doyle V. Ward, Georgia Giannoukos, Dawn Ciulla, et al. 2011. “Chimeric 16S RRNA Sequence Formation and Detection in Sanger and 454-Pyrosequenced PCR Amplicons.” Genome Research 21 (3): 494–504. https://doi.org/10.1101/gr.112730.110.

Hallen-Adams, Heather E., Stephen D. Kachman, Jaehyoung Kim, Ryan M. Legge, and Inés Martínez. 2015. “Fungi Inhabiting the Healthy Human Gastrointestinal Tract: A Diverse and Dynamic Community.” Fungal Ecology 15: 9–17. https://doi.org/10.1016/j.funeco.2015.01.006.

Holmes, Elaine, Jia V. Li, Thanos Athanasiou, Hutan Ashrafian, and Jeremy K. Nicholson. 2011. “Understanding the Role of Gut Microbiome-Host Metabolic Signal Disruption in Health and Disease.” Trends in Microbiology 19 (7): 349–59. https://doi.org/10.1016/j.tim.2011.05.006.

Hopkins, M J, R Sharp, G T Macfarlane, and South Bank. 2001. “Age and Disease Related Changes in Intestinal Bacterial Populations Assessed by Cell Culture, 16S RRNA Abundance, and Community Cellular Fatty Acid Profiles,” 198–205.

Kanehisa, Minoru, Susumu Goto, Yoko Sato, Masayuki Kawashima, Miho Furumichi, and Mao Tanabe. 2014. “Data, Information, Knowledge and Principle: Back to Metabolism in KEGG.” Nucleic Acids Research 42 (D1): 199–205. https://doi.org/10.1093/nar/gkt1076.

Keane, Sean, Pierce Geoghegan, Pedro Povoa, Saad Nseir, Alejandro Rodriguez, and Ignacio Martin-Loeches. 2018. “Systematic Review on the First Line Treatment of Amphotericin B in Critically Ill Adults with Candidemia or Invasive Candidiasis.” Expert Review of Anti-Infective Therapy 16 (11): 839–47. https://doi.org/10.1080/14787210.2018.1528872.

Llewellyn, Amy, and Andrew Foey. 2017. Probiotic Modulation of Innate Cell Pathogen Sensing and Signaling Events. Nutrients. Vol. 9. https://doi.org/10.3390/nu9101156.

Magoč, Tanja, and Steven L. Salzberg. 2011. “FLASH: Fast Length Adjustment of Short Reads to Improve Genome Assemblies.” Bioinformatics 27 (21): 2957–63. https://doi.org/10.1093/bioinformatics/btr507.

Matsubara, Victor H., H. M.H.N. Bandara, Marcia P.A. Mayer, and Lakshman P. Samaranayake. 2016. “Probiotics as Antifungals in Mucosal Candidiasis.” Clinical Infectious Diseases 62 (9): 1143–53. https://doi.org/10.1093/cid/ciw038.

Pan, Hudan, Ruijin Guo, Jie Zhu, Qi Wang, Yanmei Ju, Ying Xie, Yanfang Zheng, et al. 2018. “A Gene Catalogue of the Sprague-Dawley Rat Gut Metagenome.” GigaScience 7 (5): 1–8. https://doi.org/10.1093/gigascience/giy055.

Qiu, Xinyun, Xia Li, Zhe Wu, Feng Zhang, Ning Wang, Na Wu, Xi Yang, and Yulan Liu. 2016. “Fungal-Bacterial Interactions in Mice with Dextran Sulfate Sodium (DSS)-Induced Acute and Chronic Colitis.” RSC Advances 6 (70): 65995–6. https://doi.org/10.1039/c6ra03869g.

Qiu, Xinyun, Feng Zhang, Xi Yang, Na Wu, Weiwei Jiang, Xia Li, Xiaoxue Li, and Yulan Liu. 2015. “Changes in the Composition of Intestinal Fungi and Their Role in Mice with Dextran Sulfate Sodium-Induced Colitis.” Scientific Reports 5 (May): 1–12. https://doi.org/10.1038/srep10416.

Saulnier, Delphine M., Filipe Santos, Stefan Roos, Toni Ann Mistretta, Jennifer K. Spinler, Douwe Molenaar, Bas Teusink, and James Versalovic. 2011. “Exploring Metabolic Pathway Reconstruction and Genome-Wide Expression Profiling in Lactobacillus Reuteri to Define Functional Probiotic Features.” PLoS ONE 6 (4). https://doi.org/10.1371/journal.pone.0018783.

Seneviratne, C J, L P Samaranayake, T Ohshima, N Maeda, and L J Jin. 2016. “Identification of Antifungal Molecules from Novel Probiotic Lactobacillus Bacteria for Control of Candida Infection.” Hong Kong Medical Journal = Xianggang Yi Xue Za Zhi.

Shetty, Sudarshan Anand, Nachiket Prakash Marathe, and Yogesh S Shouche. 2013. “Opportunities and Challenges for Gut Microbiome Studies in the Indian Population,” 1–12.

Strollo, Sara, Michail S Lionakis, Jennifer Adjemian, Claudia A Steiner, and D Rebecca Prevots. 2017. “Epidemiology of Hospitalizations Associated with Invasive.” Emerging Infectious Diseases 23 (1): 7–13.

Thomas, Carissa M., and James Versalovic. 2010. “Probiotic-Host Communication: Modulation of Host Signaling Pathways.” Gut Microbes 1 (3): 148–63. www.landesbioscience.com/journals/gutmicrobes/article/11712.

Turnbaugh, Peter J, Vanessa K Ridaura, Jeremiah J Faith, Federico E Rey, and Jeffrey I Gordon. 2010. “Metagenomic Analysis in Humanized Gnotobiotic Mice” 1 (39857): 1–19. https://doi.org/10.1126/scitranslmed.3000322.

Wang, Jun, Junjie Qin, Yingrui Li, Zhiming Cai, Shenghui Li, Jianfeng Zhu, Fan Zhang, et al. 2012. “A Metagenome-Wide Association Study of Gut Microbiota in Type 2 Diabetes.” Nature 490 (7418): 55–60. https://doi.org/10.1038/nature11450.

Wang, Qiong, George M. Garrity, James M. Tiedje, and James R. Cole. 2007. “Naïve Bayesian Classifier for Rapid Assignment of RRNA Sequences into the New Bacterial Taxonomy.” Applied and Environmental Microbiology 73 (16): 5261–67. https://doi.org/10.1128/AEM.00062-07.

Wheeler, Matthew L., Jose J. Limon, Agnieszka S. Bar, Christian A. Leal, Matthew Gargus, Jie Tang, Jordan Brown, et al. 2016. “Immunological Consequences of Intestinal Fungal Dysbiosis.” Cell Host and Microbe 19 (6): 865–73. https://doi.org/10.1016/j.chom.2016.05.003.

Wong, Sarah Sze Wah, Richard Yi Tsun Kao, Kwok Yong Yuen, Yu Wang, Dan Yang, Lakshman Perera Samaranayake, and Chaminda Jayampath Seneviratne. 2014. “In Vitro and in Vivo Activity of a Novel Antifungal Small Molecule against Candida Infections.” PLoS ONE 9 (1). https://doi.org/10.1371/journal.pone.0085836.

Yuying Liu, Xiangjun Tian, Baokun He, Thomas K. Hoang, Christopher M. Taylor, Eugene Blanchard, Jasmin Freeborn, Sinyoung Park, Meng Luo, Jacob Couturier, Dat Q. Tran, Stefan Roos, Guoyao Wu, and J. Marc Rhoads. 2016. “Lactobacillus Reuteri DSM 17938 Feeding of Healthy Newborn Mice Regulates Immune Responses While Modulating Gut Microbiota and Boosting Beneficial Metabolites.” American Journal of Physisology-Gastrointestial and Liver Physiology 1607 (1): 243–243. https://doi.org/10.1007/s40278-016-18817-x.

Zangl, Isabella, Ildiko-Julia Pap, Christoph Aspöck, and Christoph Schüller. 2020. “The Role of Lactobacillus Species in the Control of Candida via Biotrophic Interactions.” Microbial Cell 7 (1): 1–14. https://doi.org/10.15698/mic2020.01.702.

