## Supplementary material for "Time-dependent modulation of gut microbiome in response to systemic antifungal agents": Supplimentary figures

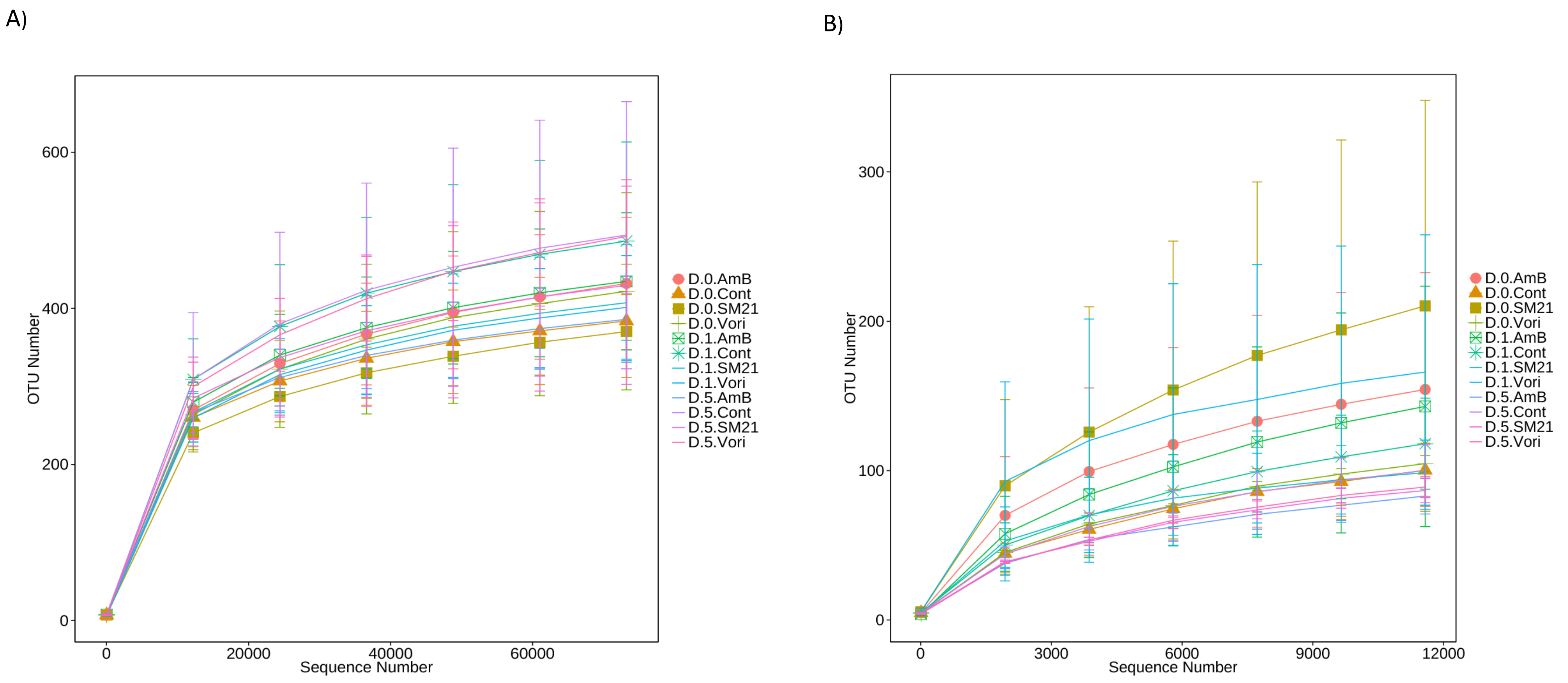


**Supplementary Figure 1:** The rarefaction curves of OTUs

The x-axis shows the number of valid sequences per sample and the y-axis shows the observed species (OTUs). Each curve in the graph represents a different sample and is shown in a different colour. As the sequencing depth increased, the number of OTUs also increased. Eventually the curves began to plateau, indicating that as the number of extracted sequences increased, the number of OTUs detected was decreased. rarefaction curves by (A) 16S rDNA region (B) ITS2 region. Although the rarefaction curves from both 16SrDNA and ITS2 did not achieve the even stage, the terminal slopes of them were rather low, suggesting that the sequencing detected the majority of the species

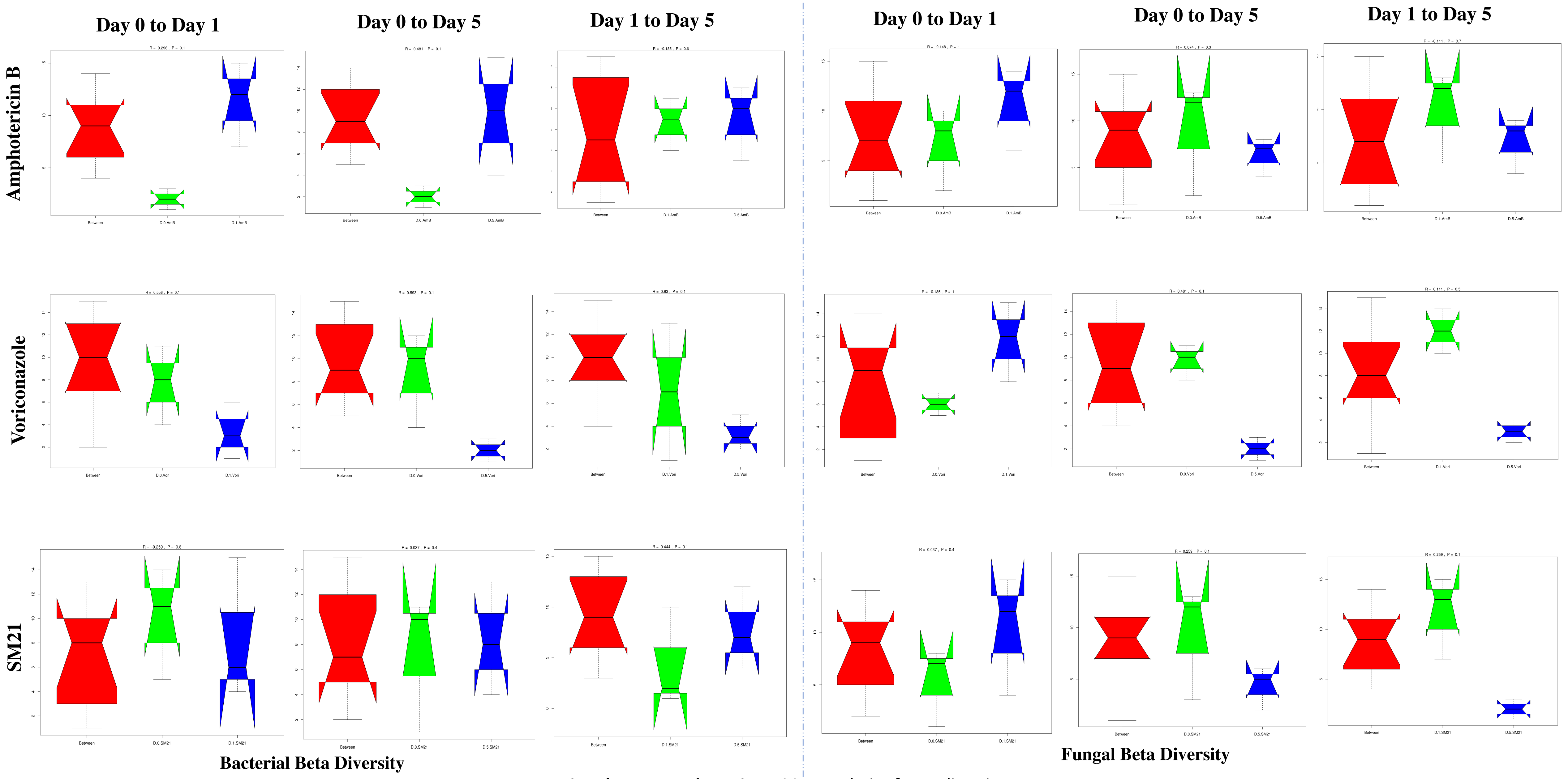


**Supplementary Figure 2:** ANOSIM analysis of Beta diversity

Box plot of Inter-group and Intra-group Beta distance (ANOSIM Analysis). The x-axis represents the grouping and the y-axis represents the distance calculated by Unweighted_unifrac. The data in the box is the distance of Inter-group and Intra-group, respectively. R-value: R-value range (–1, 1). The R-value ≤ 0 represents no significant differences of inter-group and intra-group, and R-value > 0 shows that inter-group differences are greater than intra-group differences. P-value: the P-value represents the confidence level of the statistical analysis; P < 0.05 reflects significant differences between Inter-group and Intra-group. Significant condition associated clustering was not observed in ANOSIM similarities matrices (R > 0, $P_{anosim}$ > 0.05) of any antifungal treatment.

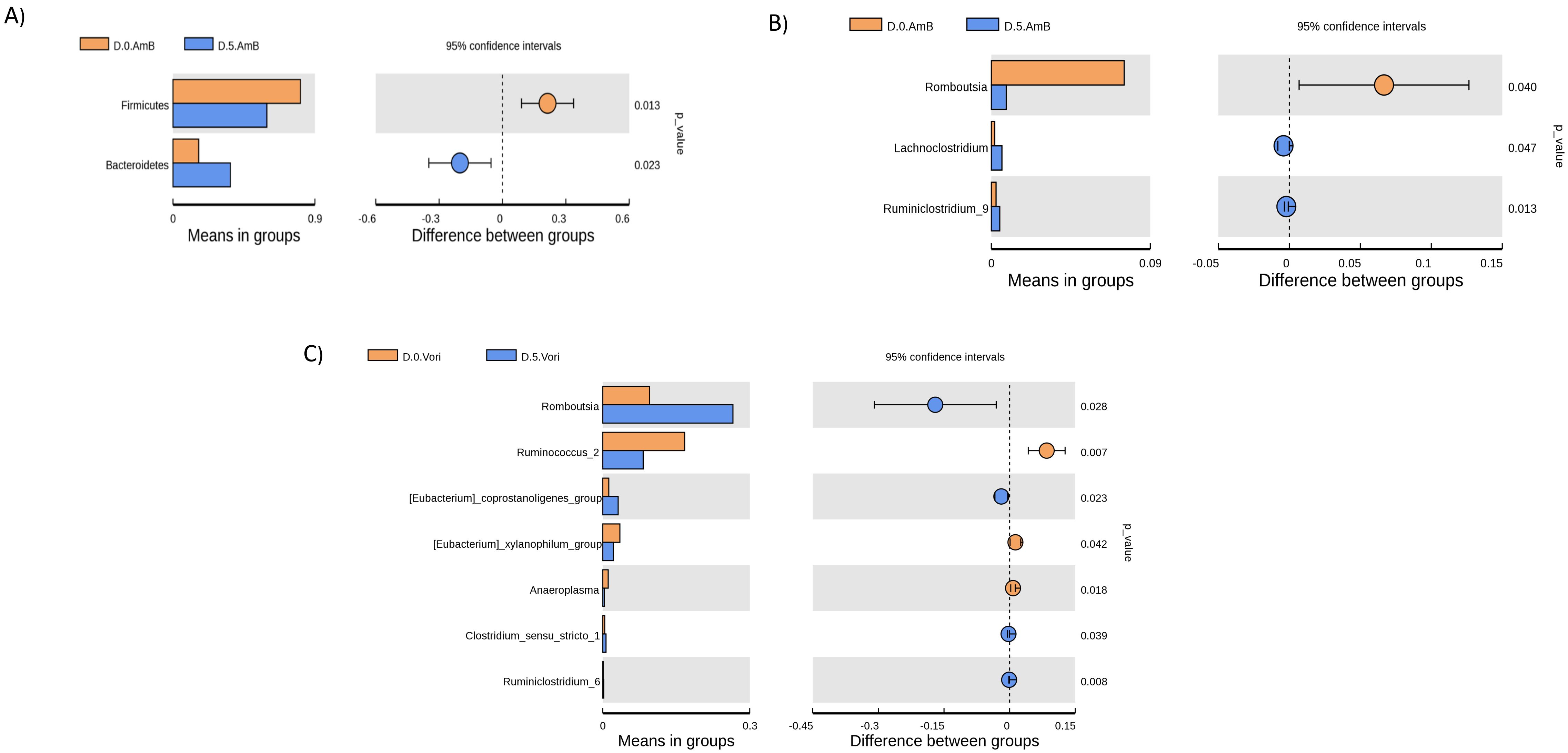


**Supplementary Figure 3:** Between-group variation analysis of species

T-test is performed to determine species with significant variation between groups (p value < 0.05) at various taxon ranks including phylum, class, order, family, genus, and species. The left panel is the abundance of species showing significant difference between group. Each bar represents the mean value of the abundance in each group of the species showing significant difference between group. The right panel is the confidential interval of between group variation. The left-most part of each circle stands for the lower limit of 95% confidential interval, while the right-most part is the upper limit. The center of the circle stands for the difference of the mean value. The colour of the circle is in agree with the group whose mean value is higher. The right-most value is the p-value of the significance test of between group variation. Significant difference between group of A) Amphotericin B(AmB) on Phylum level, B) AmB on genus level and, C) Voriconazole (Vori) on genus level. Both AmB and Voriconazole treatment dominated cluster is defined by the shared presence of microbial taxon belonging to the genus Romboustia (P= 0.028 and P= 0.040; respectively) in contrast to respective control groups.

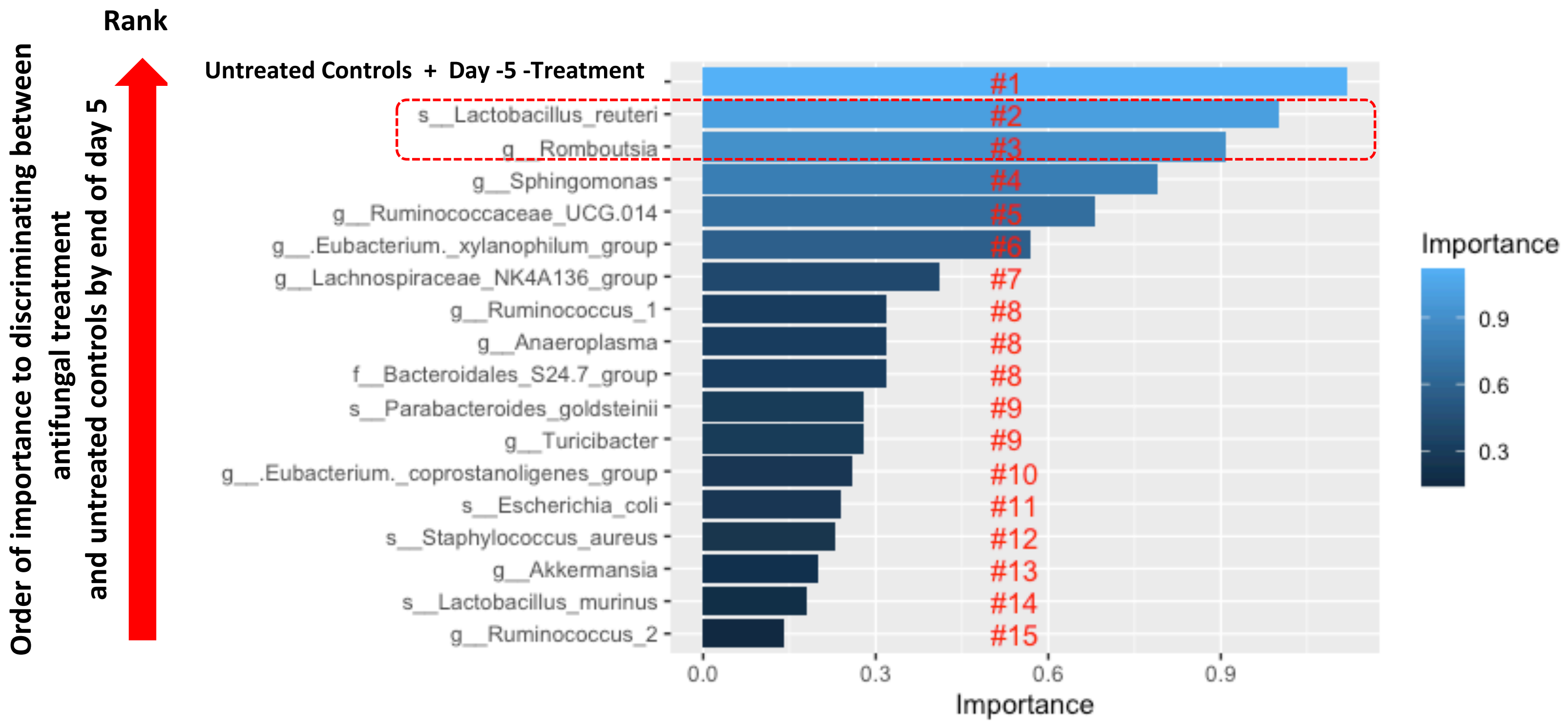


**Supplementary Figure S4.** Random Forest classification

Random Forest classification was performed on the relative abundances of 17 bacterial genera with significant and uniform differential abundances from LeFse output. 10-fold cross-validation was performed to search to select the optimal subset of bacteria discriminating among all four groups. The final model accuracy is taken as the mean from the number of repeats. Taxa within the red rectangular box are the one delineating the clustering of most important bacterial composition between control and treatment groups by end of treatment day 5, marked by highest mean importance in accuracy (OOB estimate of error rate: 41.2%).
